# Single Cell Raman-Deuterium Isotope Probing for Drug Resistance of *Elizabethkingia* spp

**DOI:** 10.1101/2021.06.08.447646

**Authors:** Shuying Yuan, Yanwen Chen, Yizhi Song, Lin Zou, Kaicheng Lin, Xinrong Lu, Ruijie Liu, Shaoxing Zhang, Danfeng Shen, Zhenju Song, Chaoyang Tong, Li Chen, Guiqin Sun

## Abstract

Nosocomial infection associated with *Elizabethkingia* spp. is an emerging clinical concern characterized by multi-drug resistance and severe clinical consequences particularly in immunocompromised individuals and infants. Efficient control of this infection demands quick and reliable methods to determine the right drugs for the treatment. In this study, *E. meningoseptica* ATCC 13253 and four clinical isolates of *Elizabethkingia* spp. obtained from China, were subjected to single cell Raman spectroscopy analysis coupling with deuterium probing (single cell Raman-DIP). The results demonstrated that single cell Raman-DIP could generate an antimicrobial susceptibility testing result for *Elizabethkingia* spp. colonies within 4 hours based on their metabolisms variations at single cell level, and the drug resistant spectra of *Elizabethkingia* spp. determined by single cell Raman-DIP were consistent with the classical MIC method. Meanwhile single cell Raman spectroscopy (single cell RS) was applied to analyze Raman spectra of *Elizabethkingia* spp., which were revealed that their ratios of nucleic acid/protein were lower than other gram-negative pathogens and isolates from different origins could be distinguished by their Raman fingerprint. The *in vitro* results confirmed that minocycline and levofloxacin are first-line antimicrobials for *Elizabethkingia* spp. infection.

## Introduction

The emergence and widespread distribution of antimicrobial resistant bacteria has led to an increasing concern with respect to potential environmental and public health risks. The crisis of antimicrobial resistant bacteria has been attributed to the overuse and misuse for medication (1). Therefore, it is important for clinicians to know the drug resistance of pathogens and use the suitable antimicrobials. In clinical practice, the bacterial drug resistance has been relied on phenotypic AST approaches, such as “minimum inhibitory concentration” (MIC) detected by broth microdilution method (BMD) (2, 3). At least 16-18 hours were taken by this method to detect the antimicrobial effects on bacterial population growth for isolated colonies. Rapid detection of microbial antimicrobial susceptibility could ensure the selection of effective antimicrobials and provide a reduction in total antimicrobial consumption (4).

Raman spectroscopy is a label-free, fast and non-destructive biochemical phenotype technology that been supplied to detect the vibration modes of molecules (5). Single cell RS provides a biochemical “fingerprint” of individual cell, which were reflected cell physiological and metabolic states (6). It was applied to identify bacterial strains (7–10) and to detect physiological changes during the treatment of antimicrobials (11, 12). It has been reported that different Raman spectra were consisted in *Listeria monocytogenes* with different susceptibilities to sakacin P in 2006 (13). Single cell RS analysis on different pathogens, including methicillin-resistant *Staphylococcus aureus* (MRSA), *Enterococcus faecium, Enterococcus faecalis, Escherichia coli, Klebsiella pneumoniae, Pseudomonas aeruginosa, Acinetobacter baumannii, Serratia marcescens* and *Lactococcus lactis*, have been carried out (14–20).

It has been discovered that the metabolically active microorganisms could incorporate the deuterium (D) into the cells via NADH/NADPH electron transport chain, producing a newly formed carbon-deuterium (C-D) band in single cell RS analysis coupling with deuterium probing (single cell Raman-DIP) (5, 6, 21, 22). The occurrence of C-D band around 2040-2300 cm^−1^ has been recognized as antimicrobial resistance biomarker when bacteria were exposed to antimicrobials and D_2_O (11). Recently, single cell Raman-DIP was proposed to achieve fast AST for pathogens such as *Escherichia coli*, *Klebsiella pneumoniae*, *Streptococcus mutans, Lactobacillus fermentum, Enterococcus faecalis and Staphylococcus aureus* (5, 23–26). However, the potential of single cell Raman-DIP application for fast AST for *Elizabethkingia* spp. has been undemonstrated.

Hospital infection associated with *Elizabethkingia* spp. is an emerging clinical concern characterized by multi-drug resistance and severe clinical consequences. Many cases of *Elizabethkingia* spp. infections have been reported as part of outbreaks in the state of Wisconsin (USA), London (UK) and Mauritius (27–29). *Elizabethkingia* spp. is a group of Gram-negative, none-ferment pathogens, responsible for a panel of diseases, including meningitis, sepsis, bacteremia, pneumonia, and neutropenic fever (30–33).

In this study, the optimized single cell Raman-DIP was applied to four hospital-isolated *Elizabethkingia* spp. strains and a standard strain *Elizabethkingia meningoseptica* (EM) ATCC 13253. Unique features of *Elizabethkingia* spp. were revealed by that single cell RS, which were improved the tested might be used to differentiate *Elizabethkingia* spp. from the other pathogens. The MIC readout of *Elizabethkingia* spp. were given by single cell Raman-DIP within 4 hours and high consistent results were shown with gold standard. The possibility of applying single cell Raman-DIP for clinical diagnosis of *Elizabethkingia* spp. was demonstrated.

## Materials and Methods

### Microorganisms and growth conditions

Five *Elizabethkingia* spp. strains were used in this study, including ATCC 13253 ordered from American Type Culture Collection and four clinical isolates (FMS-007, HS-2, NB-46 and TZ-3) collected in China (34, 35). All strains were grown aerobically in Trypticase Soy Broth (TSB) (Sigma-Aldrich, America) at 35°C overnight and then seeded on blood agar plate (BioMérieux, France) for single colonies. The tested strains were confirmed by automated VITEK 2 Compact system with the Gram-Negative identification card (GN) (BioMérieux, France). The AST results were determined by single cell Raman-DIP and MIC, respectively.

### Single cell RS analysis of *Elizabethkingia* spp

Single colonies on blood agar plate were inoculated into 1 ml TSB and incubated at 35°C with 180 rpm for 16 h. Cells were washed with sterile deionized water three times and resuspended in 10 ml sterile deionized water. After washing and resuspending, 2.5 μL of cell suspension were transferred onto an aluminum-coated slide (Sonopore, China). Raman spectra were obtained using WITec Alpha300R confocal Raman microscope (WITec, Germany) with 532 nm excitations laser, 100× magnifying dry objective (NA =0.9) (Carl Zeiss, Germany), 600 gr/mm grating. The integration time per spectrum was 20 s and the power on the sample was 7-9 mW. Spectrometry was measured for single cells spanning the range of 300-1,900 cm^−1^ to cover the most relevant Raman peaks of microbial cells. The spectra of 5 strains were analyzed with Principal component analysis (PCA) and linear discriminant analysis (LDA) method in R software using FactoMineR package as described previously (24). PCA was used to compress the information held by the spectra and the first 20 principal components that described the greatest variance of the spectral data were used for LDA.

### MIC of *Elizabethkingia* spp. determined by broth microdilution method

A standard assay (CLSI, 2019) was conducted in a 96-well microplate (Bio-kont, China). The tested bacterial colonies on each blood agar plate were aseptically transferred into Mueller-Hinton (MH) broth (BD Biosciences, America), and a homogenous suspension with a density equivalent to a 0.5 McFarland’s standard was prepared. Then the bacterial suspension was diluted with MH broth at a ratio of 1:200. One hundred microliters of prepared bacterial solution were inoculated into wells with sequentially diluted antimicrobials and incubated at 35°C with 180 rpm for 16 h. Aztreonam, cefepime, imipenem, ticarcillin/clavulanic acid, piperacillin/tazobactam, amikacin, tobramycin, minocycline, levofloxacin and trimethoprim/sulfamethoxazole were used in this study. The concentrations of the ten antimicrobials were based on the breakpoints of other Non-Enterobacteriaceae listed in guidelines M100 (CLSI, 2019). MIC value and the results of antimicrobial susceptibility were interpreted based on guidelines M100 (CLSI, 2019).

### Antimicrobial susceptibility detected by single cell Raman-DIP

D_2_O (99% D atom, Sigma-Aldrich, US) labelling was performed following previous reported studies (6, 21). 3~5 isolated colonies selected from each blood agar plate were subjected to single cell Raman-DIP. Bacterial suspension preparation was prepared as the MIC detection. One hundred microliters of inoculated bacterial solution were added to each 96-well plates with standard concentrations of antimicrobial suggested in CLSI and incubated at 35°C with 180 rpm for 1 h. Sixty-six microliters 66 μL of D_2_O was added to each well, and then incubated for two more hours. Cells were washed with sterile deionized water three times and resuspended in 50 μL of sterile deionized water. Two point five microliters of cell suspension were transferred onto an aluminum-coated slide. At least 20 to 30 single cell Raman spectra were obtained with 4 s integration time for each treatment. The Raman spectra for carbon-deuterium (C-D) peaks (2040–2300 cm^−1^) and carbon-hydrogen (C-H) peaks (2800–3100 cm^−1^) were obtained respectively. The C-D ratio was calculated (C-D/C-D+C-H) and normalized (C-D ratio of treated group minus C-D ratio of no deuterium and no antimicrobials controls). The impact of a treatment was decided by the relative metabolic rate (relative C-D rate of the treatment was calculated by the ratio of normalized C-D of the treatment and the no drug control) (24). The cutoff reads for relative metabolic rate is 0.6 to separate the metabolism active (>0.6) and inhibited conditions (<0.6) for bacteria cultured under antimicrobial treatment (26).

## Results

### Single cell Raman spectral of *Elizabethkingia* spp

The *Elizabethkingia* spp. strains used in this study were confirmed by biochemical characterization via automatic VITEK 2 Compact bacterial identification and drug sensitivity analysis system (data not shown). Raman spectra of five *Elizabethkingia* spp. strains and four ATCC Gram-negative reference strains (*Acinetobacter baumannii* ATCC 19606, *Escherichia coli* ATCC 25922, *Pseudomonas aeruginosa* ATCC 27853 and *Klebsiella pneumoniae* ATCC 700603) were measured and plotted in Figure 1A. Comparing with the other gram-negative strains, lower peaks in Raman spectra at around 780, 1479 and 1578 cm^−1^ were shown in *Elizabethkingia* spp. strains significantly. Raman peak at around 1450 cm^−1^ (assigned to protein) was chosen as a reference(36). The ratio of the peak intensity at 780, 1479 and 1578 cm^−1^ to 1450 cm^−1^ were calculated and plotted in Fig 1B, 1C and 1D, respectively. The position of a Raman peak is represented its molecular structure while the intensity of the peaks is associated with the content of the molecule. Raman peaks at 780 cm^−1^ was assigned to cytosine and uracil structure, 1479 cm^−1^ was to guanine and adenine structure, 1578 cm^−1^ was assigned to guanine and adenine structure (37–39). The results of single cell RS implied that the ratio of nucleic acid/protein in *Elizabethkingia* spp. were significantly lower than the other four Gram-negative reference strains (Fig. 1).

**Figure 1.**
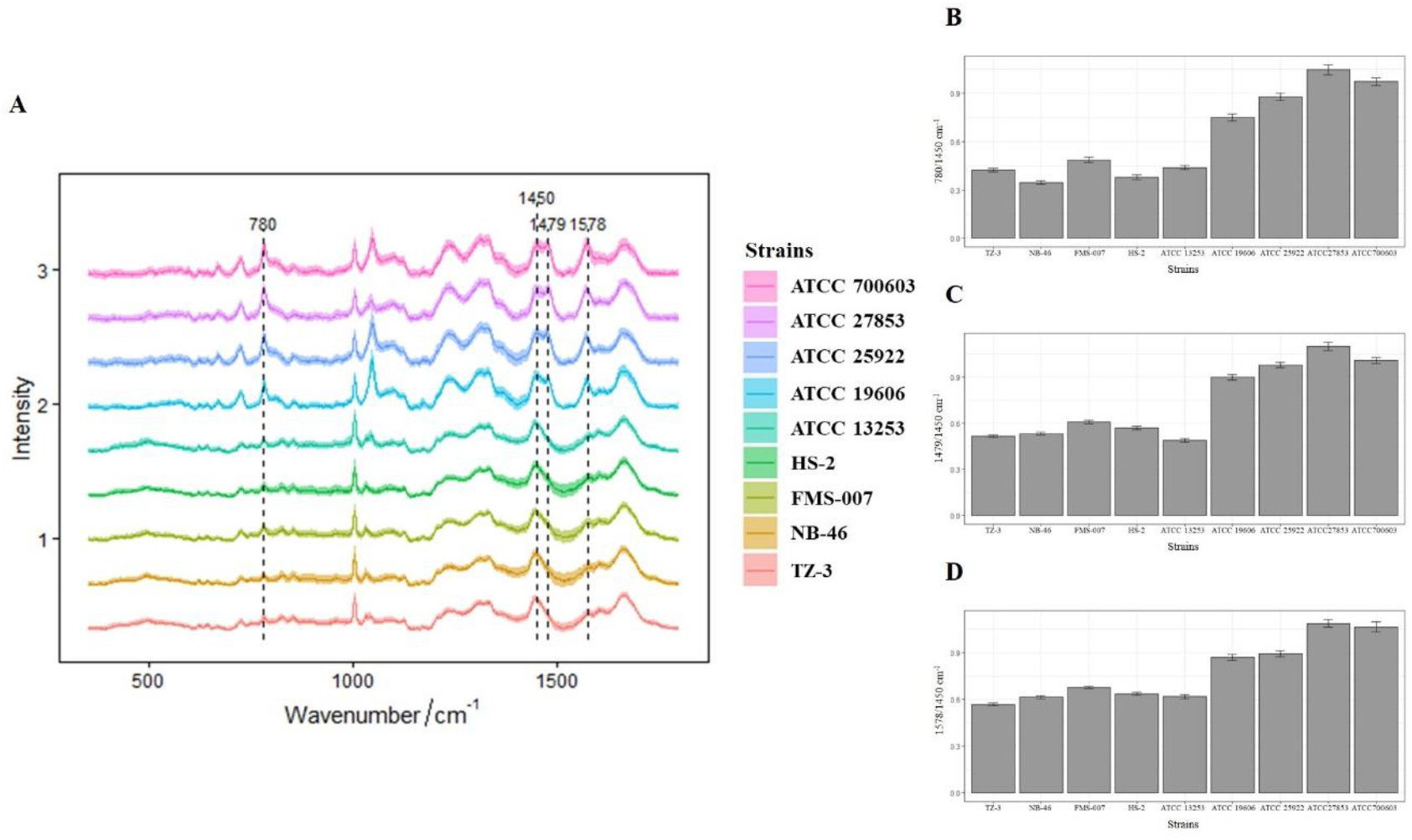
(A) Representative Raman spectra of five *Elizabethkingia* spp. strains and other four Gram-negative bacteria. (B) The ratio of nucleic acid peak at 780 cm^−1^ and protein peak at 1450 cm^−1^. (C) The ratio of nucleic acid peak at 1479 cm^−1^ and protein peak at 1450 cm^−1^. (D) The ratio of nucleic acid peak at 1578 cm^−1^ and protein peak at 1450 cm^−1^.

### Discrimination of *Elizabethkingia* spp. by single cell RS

Single cell RS of five *Elizabethkingia* spp. strains were analyzed through machine learning model LDA. The score of each spectrum on two most prominent components were illustrated in Fig. 2. It is clearly shown that the RS of five *Elizabethkingia* spp. strains were divided into two groups along the LD1 axis. One group mainly contained spectra from ATCC 13253, and the other group contained spectra from clinical strains (Fig. 2). ATCC 13253 was firstly isolated at Massachusetts in 1949 (40, 41), and the other four strains were recently isolated from China. It was indicated that single cell RS might provide a novel method for typing of *Elizabethkingia* spp. strains.

**Figure 2.**
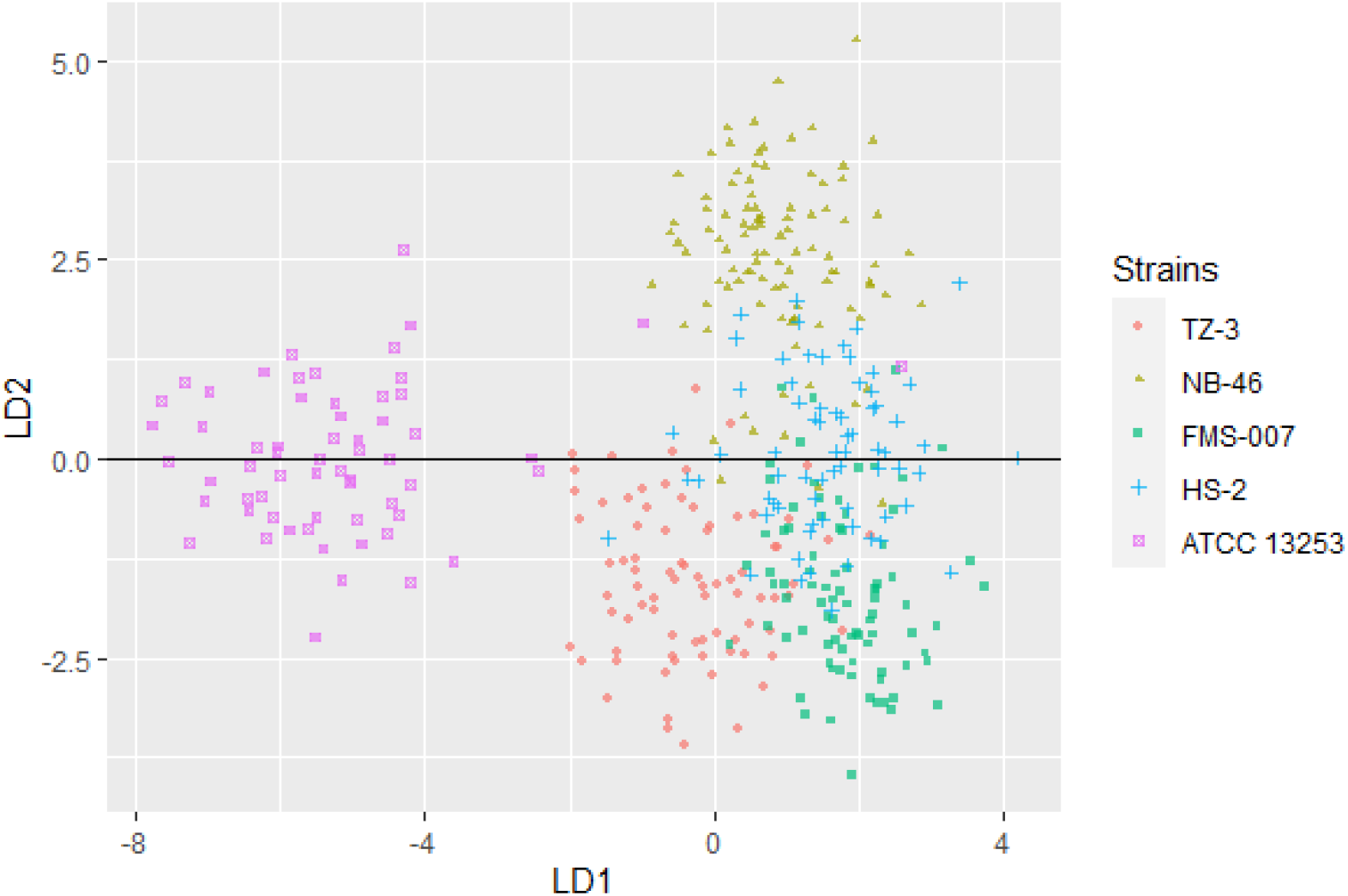
Two types of the five *Elizabethkingia* spp. strains. Each dot represented the combinative data of a single cell collected by single cell RS.

### Rapid AST of Imipenem for *Elizabethkingia* spp. by single-cell metabolic activity

Single cell Raman-DIP was applied to measure metabolic activity of bacteria at single cell level (21). *Elizabethkingia* spp. strains were incubated in MH medium containing 40% of heavy water for 2 hours, the deuterium in the medium was incorporated in the cellular biomass and produced a new peak at 2040-2300 cm^−1^ (C-D band) on the basis of 2800-3100 cm^−1^ (C-H band) (Fig. 3A and 3D). Taken imipenem as an example to detect metabolism under antimicrobials treatment using single cell Raman-DIP. After exposed in imipenem, the intensity of C-D band in Strain FMS-007 was remained at the same level of the drug-free control in which no imipenem was added (Fig. 3A), then C-D ratio (Fig. 3B) and normalized C-D ratio (Fig. 3C) were calculated. This was indicated that bacteria were still metabolic active under imipenem up to 16 ug/ml and resulted in an MIC of above 16 ug/ml. Since 16 ug/ml was the resistant breakpoint of *Elizabethkingia* spp. for imipenem, strain FMS-007 was regarded resistant to imipenem. The C-D band of strain TZ-3 was not detected when imipenem was presented at a concentration of 16 μg/ml (Fig. 3D, 3E and 3F), which meant the bacterial metabolic activity was inhibited and the MIC could be determined as 16 ug/ml. The results showed that single cell Raman-DIP could be used to detect microbial metabiotic activity and antimicrobial susceptibility of *Elizabethkingia* spp.

**Figure 3.**
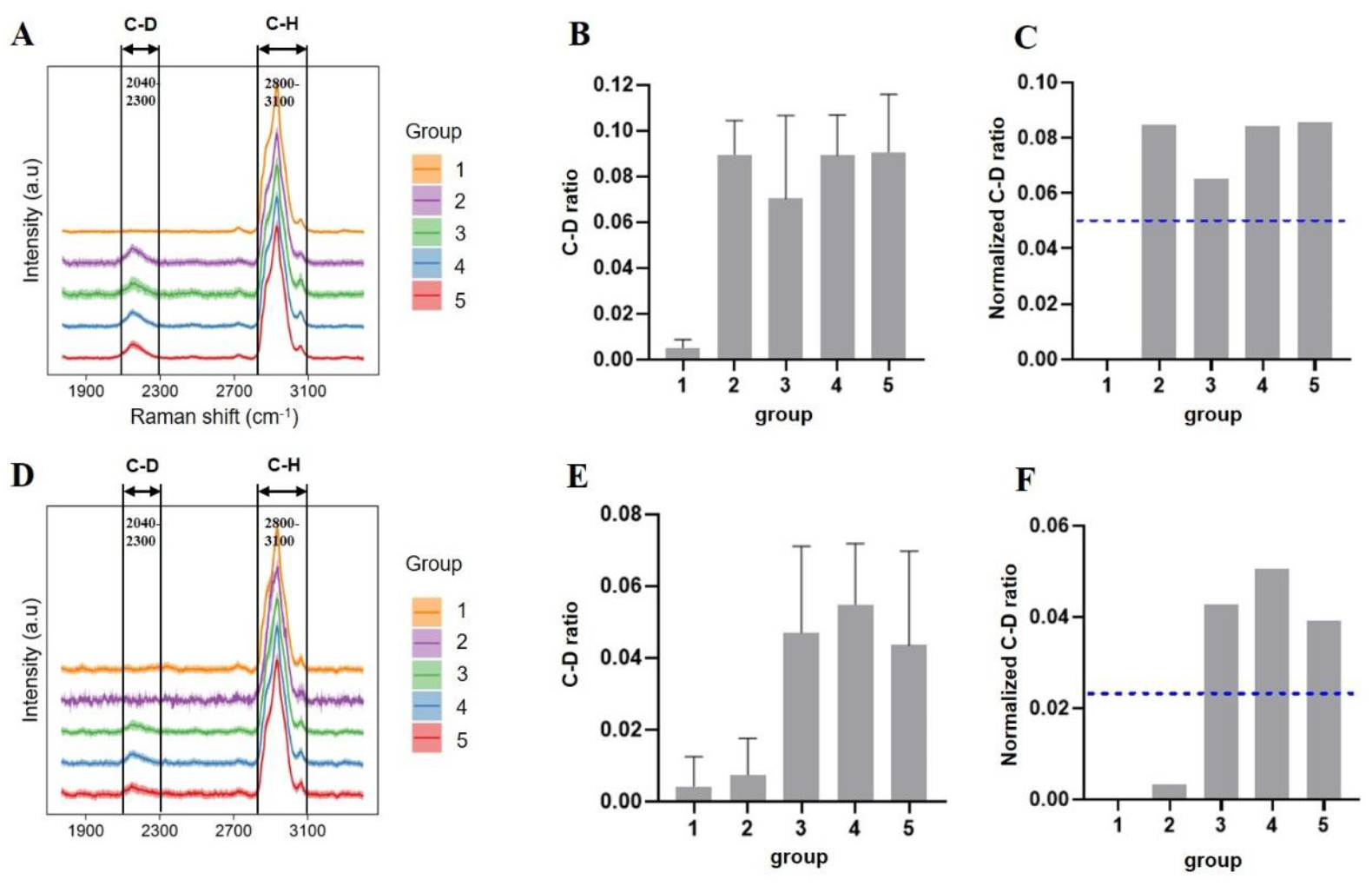
Raman spectra of cells treated with imipenem. (A) C-D and C-H band in FMS-007. (B) C-D ratio of FMS-007. (C) Normalized C-D ratio of FMS-007. (D) C-D and C-H band in TZ-3. (E) C-D ratio of TZ-3. (F) Normalized C-D ratio of TZ-3. The dotted lines in (B), (C), (E) and (F) indicate the cutoff value at 0.6 of the normalized C-D ratios. Group 1: without D_2_O and imipenem; Group 2: D_2_O with 16 ug/ml imipenem; Group 3: D_2_O with 8 ug/ml imipenem; Group 4: D_2_O with 4 ug/ml imipenem; Group 5: D_2_O without imipenem.

### The AST of *Elizabethkingia* spp. by single cell Raman-DIP

The antimicrobial susceptibility of five *Elizabethkingia* spp. strains to ten antimicrobials were tested by single cell Raman-DIP and classical MIC method. The results by single cell Raman-DIP and MIC were summarized in Figure 4 and Table 1, respectively. The breakpoints of antimicrobial susceptibility on guidelines M100 (CLSI, 2019) were used in this study.

**Figure 4.**
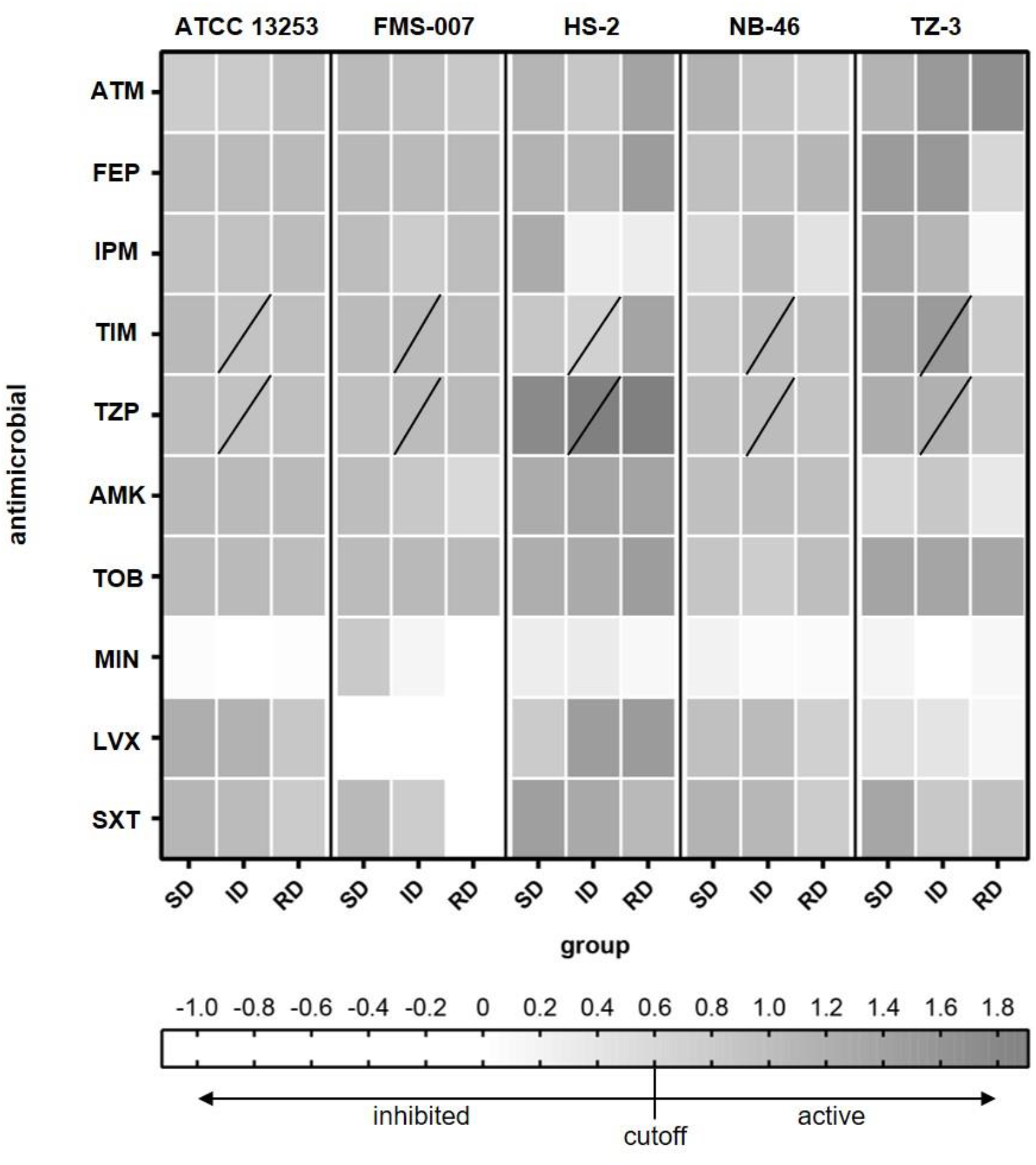
The relative metabolic rate of the five *Elizabethkingia* spp. strains determined by Raman-DIP. The antimicrobial concentrations of group SD, ID and RD were referenced by guidelines M100 (CLSI, 2019). SD: susceptible breakpoint corresponds antimicrobial concentration; ID: intermediate breakpoint corresponds antimicrobial concentration; RD: resistant breakpoint corresponds antimicrobial concentration; ATM: aztreonam; FEP: cefepime; IPM: imipenem; TIM: ticarcillin/clavulanic acid; TZP: piperacillin/tazobactam; AMK: amikacin; TOB: tobramycin; MIN: minocycline; LVX: levofloxacin; SXT: sulfamethoxazole/trimethoprim. TIM and TZP were composed of two intermediate breakpoints (the data of lower concentration of intermediate breakpoints was shown);

**Table 1.**
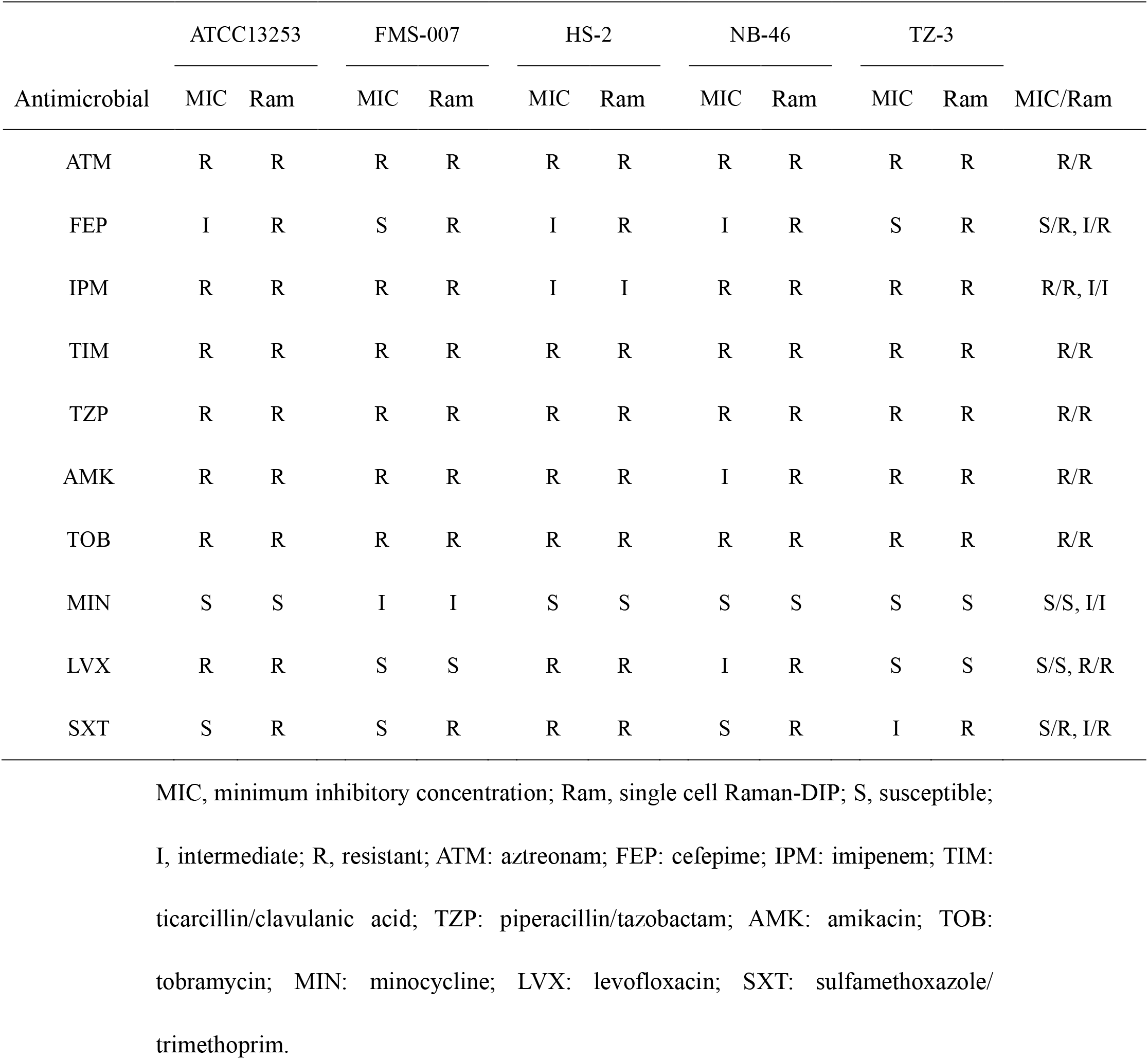
The antimicrobial susceptibility of five *Elizabethkingia* spp. strains

The corresponding concentration of relative metabolic rate <0.6 was used to interpreted the result of antimicrobial susceptibility. In the AST results tested by single cell Raman-DIP, all five strains were resistant to seven antimicrobials (total tested ten antimicrobials), including aztreonam, cefepime, ticarcillin/clavulanic acid, piperacillin/tazobactam, amikacin, tobramycin and sulfamethoxazole/trimethoprim (Fig. 4). Except strain HS-2 was intermediate, they were resistant to imipenem. ATCC13253. HS-2, NB-46 and TZ-3 were sensitive to minocycline, while FMS-007 was intermediate. FMS-007 and TZ-3 was sensitive to levofloxacin, but ATCC 13253, HS-2 and TZ-3 were resistant. It implied that minocycline and levofloxacin were selectable drugs for *Elizabethkingia* spp. infection.

The antimicrobial susceptibility of five *Elizabethkingia* spp. strains were also tested by MIC, and the AST results between MIC and single cell Raman-DIP were compared (Table 1). The results of eight antimicrobials were consistent detected by single cell Raman-DIP and classical MIC method, including aztreonam, imipenem, ticarcillin/clavulanic acid, piperacillin/tazobactam, amikacin, tobramycin, minocycline and levofloxacin. However, the AST results of two methods were different in cefepime and sulfamethoxazole/trimethoprim. Five strains were all resistant to cefepime and sulfamethoxazole/trimethoprim tested by single cell Raman-DIP, but the results tested by MIC were sensitive or intermediate.

In this study, the results of antimicrobial susceptibility were similar between single cell Raman-DIP and MIC method. It took 4 hours to gain the AST results by single cell Raman-DIP, which was 4-5 times faster than MIC method. The results demonstrated that AST determined by single cell Raman-DIP was comparable to MIC, and single cell Raman-DIP was a practical complementation for MIC.

## Discussion

Building a quick and reliable drug-resistance assay is an urgent clinical need for medical practice and public health. In this study, single cell Raman-DIP analysis was applied to measure the fingerprint and metabolic profile of *Elizabethkingia* spp. to a panel of antimicrobials. The isolates of *Elizabethkingia* spp. were responded high sensitivity to minocycline and levofloxacin confirmed by 4-hour single cell Raman-DIP procedure. It was consistent with previous reports (30, 35, 42–44). Our study on *Elizabethkingia* spp. was an addition to the previous single cell Raman-DIP application.

In few cases, the inconsistent susceptibility results were presented between single cell Raman-DIP and MIC method. Apart from cefepime and sulfamethoxazole/ trimethoprim, the other eight antimicrobials were showed the overall consistency rate of single cell Raman-DIP to 95%. Notably, in all inconsistent cases, the results given by single cell Raman-DIP were resistant while were sensitive or intermediate judged by MIC method which means they were no major errors. These results met the FDA requirements (category agreement ≥90%, minor error ≤10.0%, major error ≤3.0%, very major error ≤1.5%) (26). Inconsistent results were shown in cefepime and sulfamethoxazole/trimethoprim between single cell Raman-DIP and MIC. Cefepime works by inhibiting penicillin binding proteins essential to cell wall formation (45). A possible explanation for cefepime was that the replication of bacteria under the treatment of cefepime with concentration higher than MIC was inhibited, the cells were still metabolic active. Recently, the growth-arrested bacteria may still exhibit metabolic activity has been revealed on metabolism of *Mycobacterium tuberculosis* (46) and persister bacteria treated with antimicrobials (47). Sulfamethoxazole/trimethoprim is a competitive inhibitor of dihydropteroate synthase that was involved in DNA synthesis (48). However, the reason caused by sulfamethoxazole/trimethoprim was unclear. Single cell metabolic activity assessment via single cell Raman-DIP could provide new insights on antimicrobial administration to patients.

Single cell RS has already been used to identify *Mycobacterium tuberculosis* and yeast species (11, 12). In this study, the characteristic peaks of *Elizabethkingia* spp. on single cell RS were identified (Fig. 1). Comparing to the controls, the ratio of nucleic acid/protein (the ratio of the peak intensity at 780, 1479 and 1578 cm^−1^ to 1450 cm^−1^) were significantly lower than the other gram-negative strains. The reason behind observation was required further multi-omics studies. In addition, clinical isolates of *Elizabethkingia* spp. from Chinese hospitals and reference strain ATCC 13253 isolated from Massachusetts were classified to two groups according to machine learning model on single cell RS (Fig. 2). These results were indicated that single cell RS could be used as a new and fast way for strain characterization.

The infection caused by *Elizabethkingia* spp. is a rare but life-threatening medical condition (3). A critical challenger for clinician is to prescribe a useful antimicrobial although strains are multidrug resistant. In this study, we demonstrated that single cell Raman-DIP could achieve a rapid and reliable AST of *Elizabethkingia* spp. in 4 hours. Single-cell RS combined with machine learning could distinguish *Elizabethkingia* spp. from other gram-negative pathogens. Thus, great potential in *Elizabethkingia* spp. related clinical studies and diagnosis were showed by single cell RS and single cell Raman-DIP technique.

## Acknowledgments

The authors would like to thank the funding support from the Natural Science Foundation of Zhejiang Province (Y20C050003), National Natural Science Foundation of China (31600644) and Department of Education of Zhejiang Province (FX2020021).

